# Identification of a sex-determining locus potentially involved in a conflict over sex-ratio

**DOI:** 10.1101/2025.05.14.654164

**Authors:** Baptiste Lhéraud, Yann Dussert, Mohamed Amine Chebbi, Isabelle Giraud, Richard Cordaux, Jean Peccoud

**Affiliations:** Laboratoire Écologie et Biologie des Interactions (UMR 7267 CNRS, Université de Poitiers), 3 rue Jacques Fort, 86000 Poitiers, France; laboratoire d’Écologie Microbienne (UMR 5557 CNRS, Université Claude Bernard Lyon 1; UMR 1418 INRAE, VetAgro Sup), 10 Rue Raphaël Dubois, 69622 Villeurbanne, France; Laboratoire Génomique Métabolique, Genoscope, Institut François Jacob (UMR 8030 CEA, CNRS, Université d’Évry, Université Paris-Saclay), 2 rue Gaston Crémieux, 91057 Évry, France; DNA Script SAS, 67 Av. de Fontainebleau, 94270 Le Kremlin-Bicêtre; Laboratoire Évolution Génomes Comportement Écologie (UMR 9191 Université Paris-Saclay, CNRS, IRD), 12 route 128, 91190 Gif-sur-Yvette, France

## Abstract

Sex-determining genes remain largely uncharacterized outside classical models in vertebrates and insects, leaving a gap in our understanding of their evolutionary emergence and sex chromosome formation. Terrestrial isopods, particularly the common pill bug *Armadillidium vulgare*, provide an excellent model for investigating these processes due to the rapid turnover in sex-determination mechanisms they undergo. In *A. vulgare*, multiple genetic determinants coexist. Notably, a feminizing factor is transmitted at a rate exceeding Mendelian expectations, resulting in female-biased populations. Some lineages possess a masculinizing allele at a locus referred to as the “*M* gene”, which is functionally analogous to an XY system. Its masculinizing dominant allele is hypothesized to have been selected due to the deficit of males caused by the feminizing factor. The existence of the *M* gene was inferred from crosses carried out in the 1990s, but its molecular nature remains unresolved. Here, we conducted a genome-wide SNP analysis combining pooled sequencing of male and female progenies with sequencing of individual parents across two families. Bayesian estimation of haplotype frequencies in progenies enabled us to delimit a candidate genomic region of approximately two megabases containing 34 annotated genes. Most notably, one of these genes encodes the androgenic gland hormone, a protein involved in male sexual differentiation. Our findings lay the groundwork for detailed genetic and functional investigations of the *M* gene, offering novel insights into the dynamics of sex determination in terrestrial isopods and into the turnover of sex chromosomes in response to sex-ratio distortion.

## Introduction

Although initially discovered using their morphology, not all sex chromosomes are heteromorphic (i.e., morphologically different) (Furman *et al*., 2020). Indeed, chromosomes that have recently acquired a sex-determining allele do not display major visible difference between homologs. Consequently, it may be difficult to identify, based on cytological observations alone, whether a diploid species shows male heterogamety (XY males and XX females) or female heterogamety (ZW females and ZZ males). On the other hand, such young sex chromosomes make it easier to identify the specific loci that control sex because these loci constitute the only systematic genetic differences between females and males (Kamiya *et al*., 2012; Akagi *et al*., 2019; Becking *et al*., 2019; Charlesworth, 2019). Beyond their profound influence on the phenotype and on chromosomal evolution, another interest of characterizing sex-determining loci stems from their involvement in genetic conflicts with non-mendelian sex-determining factors or selfish genetic elements distorting sex-ratios (Werren and Beukeboom, 1998; Uller *et al*., 2007; Kozielska *et al*., 2010; Abbott *et al*., 2017). To better identify and understand these conflicts, it is necessary to precisely identify the genes involved.

The common pill bug *Armadillidium vulgare* (Isopoda: Oniscidea) provides a good example of genetic conflicts over sex-ratio (Cordaux *et al*., 2011), as several sex determinants segregate in its populations. Among these, a ZW locus has recently been characterized as a short genomic region (Cordaux *et al*., 2021). Other lineages harbour *Wolbachia*, an intracellular maternally transmitted bacterium that has a feminizing effect inducing strong female biases in progenies (Bouchon *et al*., 1998). Yet another feminizing genetic factor called the *f* element (Legrand *et al*., 1984) consists of an insertion of the *Wolbachia* genome into the pill bug genome (Legrand *et al*., 1984; Leclercq *et al*., 2016) and is transmitted in a non-Mendelian fashion, often resulting in a sex-ratio biased towards females (Legrand *et al*., 1984; Cordaux and Gilbert, 2017). The last known sex-determining locus is called the *M* gene, which possesses a dominant masculinizing allele (Rigaud and Juchault, 1993), and is therefore akin to an XY locus. This masculinizing allele (hereafter called “*M*”) has an epistatic effect on both female heterogametic determinants: an individual carrying the W allele or the *f* element and the *M* allele develops into a functional male (Rigaud and Juchault, 1993). Such male may however retain female genital apertures (Rigaud and Juchault, 1993; Azzouna *et al*., 2004), which are assumed to arise from the feminizing element’s expression being counteracted by the *M* allele during sexual differentiation (Rigaud and Juchault, 1993). The *M* allele could therefore represent an intermediate state towards a more adapted, fully masculinizing element, which may indicate its recent origin.

One hypothesis to explain the emergence of the *M* allele and its maintenance in several populations is its ability to re-establish balanced sex ratios in populations in which the *f* element is present (Rigaud *et al*., 1997). This finds support in theoretical models showing that sex-ratio distorters can promote the turnover of sex chromosomes (Kozielska *et al*., 2010). To date, it is not known whether the *M* allele emerged recently or whether it was already present and increased in frequency following the appearance of the *f* element. To assess the age of the *M* allele and the mechanisms by which it masculinizes its carrier, we identified the genomic regions carrying the *M* allele and the protein-coding genes that could be affected by it, using a whole-genome comparison approach between daughters and sons of fathers carrying the *M* allele.

## Methods

### Detection of genomic regions linked to the *M* gene

Our approach, detailed in Cordaux *et al*. (2021), is based on crosses in which parents are individually sequenced, and their daughters and sons are sequenced in pools (pool-seq). The approach does not rely on major genetic differences between alleles, but on single nucleotide polymorphisms (SNPs) which, like the *M* gene, are heterozygous in the father and homozygous in the mother. We will refer to such SNP as “informative”, and to the allele carried by the father only as “paternal”. The following equation is used to estimate the number of recombination events (*n*_rec_) that have taken place between a SNP and the *M* gene during meiosis:

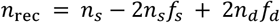

where *n*_*s*_ is the number of sons, *f*_*s*_ the frequency of the paternal allele in sons, *n*_*d*_ is the number of daughters, and *f*_*d*_ the frequency of the paternal allele in daughters. This equation assumes that the paternal allele is carried by the DNA molecule carrying the dominant sex-determining allele (here, the *M* allele), but the method also considers that the paternal allele might be carried by the homologous chromosome, a case that we will not detail here.

To compute the posterior probabilities of *f*_*s*_ and *f*_*d*_, counts of sequencing read pairs covering a given SNP and carrying each allele (paternal or alternative) are used in a Bayesian approach accounting for sequencing errors (Cordaux *et al*., 2021). *f*_*s*_ and *f*_*d*_ are averaged over all possible values weighted by their respective posterior probabilities. In practice, SNPs are not considered independently: those belonging to the same short genomic region (here a contig of a fragmented genome assembly) are assumed to be inherited identically and are combined into haplotypes. This combination drastically increases read counts, hence the precision of estimates.

Once the number of recombination events has been estimated, simulations were carried out to determine whether this number was significantly lower than expected for a contig genetically independent of the *M* gene (Cordaux *et al*., 2021). Briefly, for each contig and family, the number of read pairs carrying the paternal allele at each SNP was simulated under the assumption that the contig was unliked to the *M* gene, then *n*_rec_ was computed as detailed above. This procedure was repeated 1,000 times per contig.

### Lineages and families

To implement this approach, we used an *A. vulgare* laboratory line called “BFog”, sampled in Nice, France, in 1967. This line produces males presenting female genital apertures. Two families of the BFog line were selected. In the first family, 20 daughters and 16 sons were used and in the second family, 30 daughters and 30 sons were used. In both families, only sons with female genital apertures were selected. To verify that these males carried the W allele, and by extension the *M* allele, the presence and zygosity of the W allele were determined in the offspring and parents of the second family (there was no genetic material left for the first family) via quantitative PCR (Supplementary Text, Figure S1).

DNA was extracted from all individuals using Qiagen’s DNeasy blood and tissue kit. To produce equimolar pools of offspring, the DNA concentrations were estimated using a Qubit fluorometer (Invitrogen). Parents and offsprings were sequenced using Illumina technology (see Table S1 for information and statistics).

Reads were aligned to the *A. vulgare* reference genome (Chebbi *et al*., 2019) using BWA mem 0.7.13 (Li and Durbin, 2009). PCR duplicates were identified using samtools markdup 1.9 (Danecek *et al*., 2021). SNPs were called by the GATK algorithm using ANGSD (Korneliussen *et al*., 2014). Subsequent analyses were performed with scripts written in R (R Core Team, 2020) modified from Cordaux *et al*., (2021). These scripts select informative SNPs, estimate the frequency of paternal alleles in F1 pools hence the number of recombination events between a contig and the *M* gene, and perform the simulations described in the previous Methods section.

### Analysis of contigs linked to the *M* gene

The protein-coding gene annotation in candidate contigs was manually curated using transcriptomic and protein evidence, in case gene models were missing from the reference gene prediction. Previously published *A. vulgare* RNA-seq reads (Lewis *et al*., 2018) were trimmed with Trimmomatic 0.39 (Bolger *et al*., 2014) (parameters: ILLUMINACLIP:TruSeq3-PE-2.fa:2:30:10:8:TRUE LEADING:3 TRAILING:3 SLIDINGWINDOW:4:15 MINLEN:36), to discard adapters and low quality bases, and mapped onto the reference using STAR 2.7.1a (Dobin *et al*., 2013). For protein evidence, arthropod protein sequences from OrthoDB v11 (Kuznetsov *et al*., 2023) were aligned to the genome sequence with miniprot 0.11-r234 (Li, 2023). Manual curation of protein-coding genes was carried out using Apollo 2.7.0 (Dunn *et al*., 2019). Based on RNA-seq and protein alignments, new gene models were added when they were supported by one type of evidence, and existing models were corrected when necessary. The putative function of proteins coded by annotated genes was determined using InterProScan 5.59-91.0 (Jones *et al*., 2014) and blastP in BLAST+ 2.14.1 (Camacho *et al*., 2009) with the SwissProt 2023-06-28 database. These two analyses were carried out on the Galaxy web server at https://usegalaxy.eu/ (The Galaxy Community *et al*., 2024).

To precisely identify regions that potentially control sex, contigs identified as not having recombined with the *M* gene (*n*_rec_ < 0.5) were scanned for so-called “causal” SNPs, which are SNPs that could directly contribute to sex determination. Like the *M* gene, a SNP was considered causal if it was heterozygous in all males of both families and homozygous in all females for the same alleles. We also checked that the paternal allele was absent in *A. vulgare* lineages lacking the *M* allele, for which we had whole-genome sequence data (Table S2). Causal SNPs located in coding exons were characterized for their effect on protein sequence by a script written in R.

To check for the possibility that causal mutations could be small insertions or deletions (indels), which were not considered by our SNP-based approach, we carried out a supplementary small variant detection on the parents of both families. Variants were called using FreeBayes 1.3.1 (Garrison and Marth, 2012) imposing a minimal mapping quality of 20, a minimal base quality of 15 and no population priors. Variants of low quality or with no read support for the alternate allele on both strands and on each side of the site were then filtered out using vcffilter (QUAL > 10 & QUAL / AO > 10 & RPL > 0 & RPR > 0 & SAF > 0 & SAR > 0) in vcflib (Garrison *et al*., 2022). Variants were further filtered using vcftools 0.1.16 (Danecek *et al*., 2011) imposing a minimal sequencing depth of 5, a minimal allele count of 1 and no more than one sample with missing data. Complex variants were split with vcfallelicprimitives in vcflib. Finally, informative indels (i.e., heterozygous in the fathers and homozygous in the mothers) were selected using an R script. The genomic position of these informative indels compared to protein-coding genes was determined using the intersect function in bedtools 2.30.0 (Quinlan and Hall, 2010).

To assess whether the *M/m* polymorphism was ancient (*m* referring to the ancestral non-masculinizing allele), two analyses were carried out. First, we determined whether contigs genetically close to the *M* gene presented more SNPs than the rest of the genome, resulting from the divergence between the *M* and *m* alleles. To this aim, we tested the correlation between the density of heterozygous SNPs in the fathers with their genetic distance to the *M* gene, for contigs whose distance could be estimated with sufficient precision (paternal allele frequency estimate with a posterior probability ≥ 0.3 in every F1 pool, see Supplementary Material). Second, we tested the homology between contigs that did not recombine with the *M* gene and Y-specific contigs in *Armadillidium nasatum* (Becking *et al*., 2019), given that both systems correspond to male heterogamety. To this aim, amino-acid sequences of protein-coding genes were aligned on the Y-specific sequences of *A. nasatum* using tblastn in BLAST+ 2.12.0, keeping hits with a query coverage > 70%.

## Results

### Identification of contigs linked to the *M* gene

Genotyping of the parents using whole genome sequence data (Table S1) identified 5,555,009 informative SNPs (homozygous in the mother and heterozygous in the father) in the first family and 7,931,937 in the second family; 1,189,329 of these SNPs were common to both families. At these SNPs, computed allele frequencies in F1s allowed us to estimate the genetic distance of 39,761 contigs to the *M* gene. This resulted in the identification of 966 contigs (file S1) that recombined less with the *M* gene than expected if they were located on a different chromosome pair (Figure 1). Only six of them belonged to the 1,006 contigs attributed to the ZW chromosomes by Cordaux *et al*., (2021). As these six contigs were located between 6.5 cM and 30 cM from either the *M* or ZW locus, evidence that both sex-controlling genes are on the same chromosome pair was weak, in agreement with previous results (Rigaud and Juchault, 1993). According to the size of genomic contigs that showed significant genetic linkage to the *M* gene, the chromosome pair carrying it would measure at least 46.1 Mbp (Figure 1). By comparison, the average chromosome length of *A. vulgare* is estimated at 64 Mb (Cordaux *et al*., 2021).

**Figure 1.**
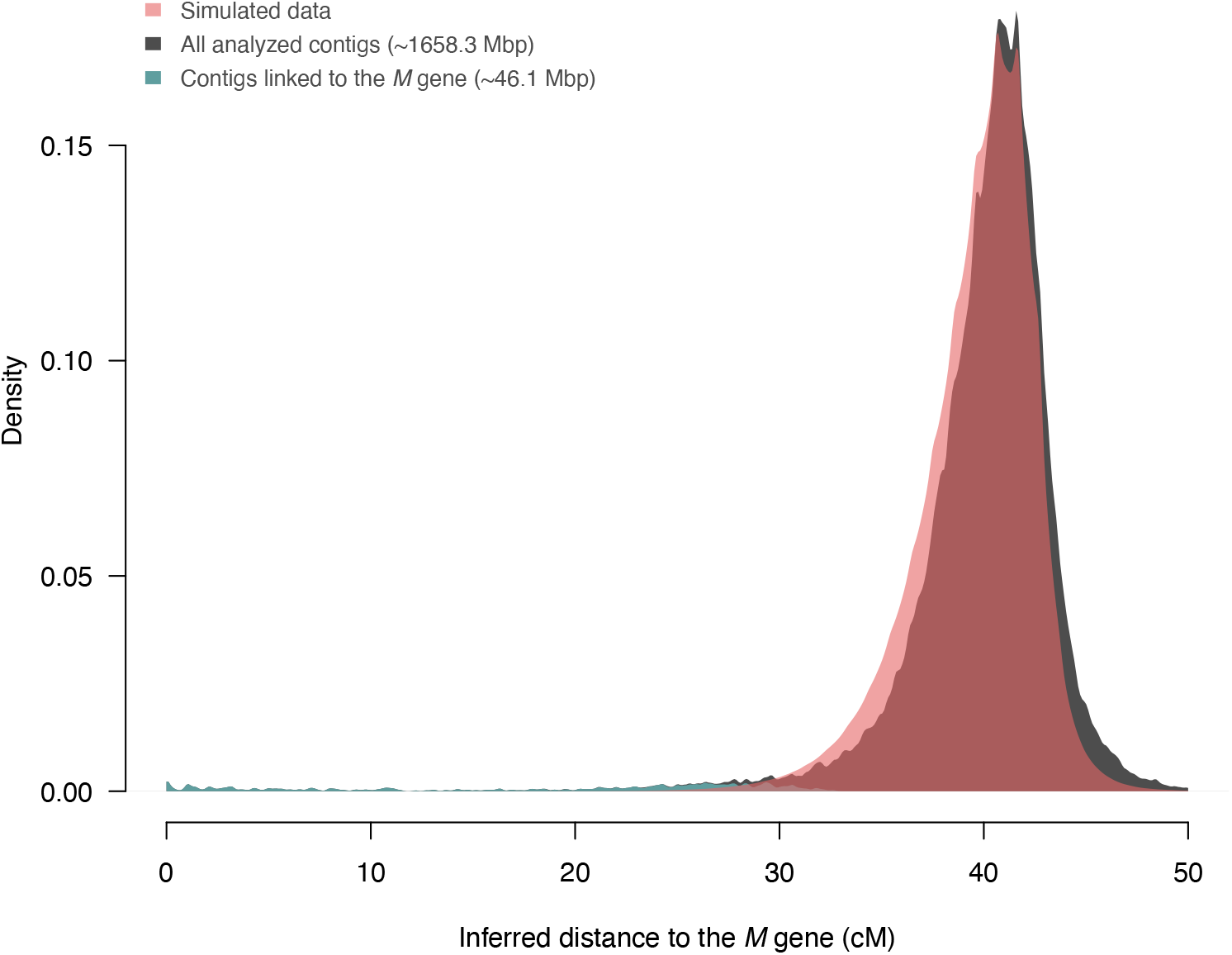
Distributions of inferred genetic distances between *Armadillidium vulgare* contigs and the *M* gene for real (black and grey-blue) and simulated data (red). Simulated data were generated under the assumption that contigs were genetically independent of the *M* gene (see Methods). Contigs linked to the *M* gene (plotted in grey-blue) are those with observed genetic distance lower than measured in all simulations.

The density of heterozygous SNPs in the fathers (Figure 2) did not increase at close distances from the *M* gene and therefore did not show evidence of an accumulation of mutations between the *M* and *m* alleles. No protein-coding gene found in the contigs linked to the *M* locus had a hit on Y-specific sequences of *A. nasatum*.

**Figure 2.**
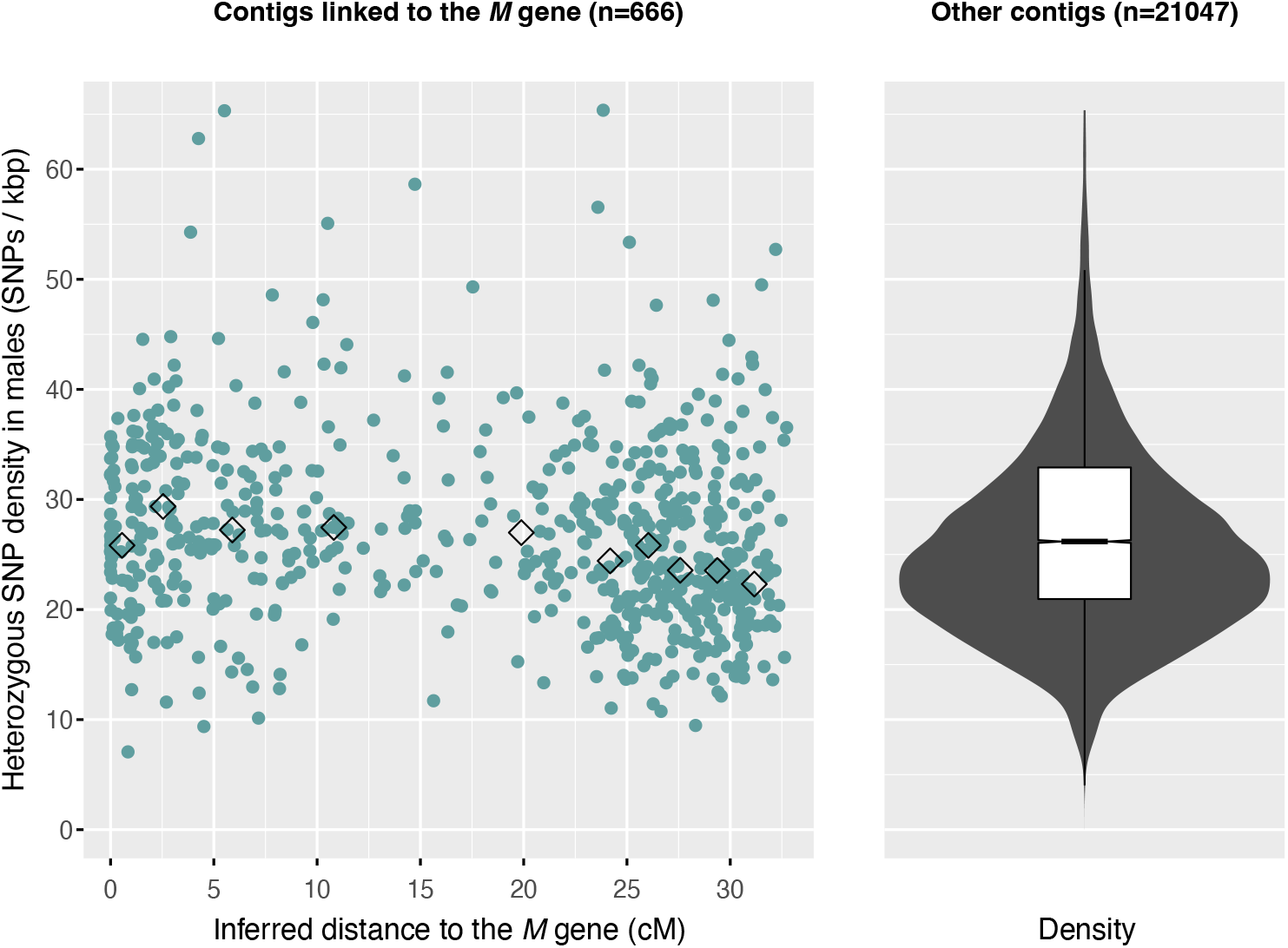
Density of heterozygous SNPs near the *M* gene. Densities of heterozygous SNPs in fathers as a function of the inferred genetic distance to the *M* gene for contigs that are statistically linked to this gene (left panel) and for other contigs (right panel). Diamonds on the left-hand plot represent averages for ten genetic distance classes delineated by deciles.

### Genetic variation that may constitute the *M* gene

Only 36 contigs (~1.9 Mb, Figure 3) did not show any signal of recombination with the *M* gene during meiosis (<0.5 inferred recombination events). They include 34 predicted genes (**Erreur ! Source du r envoi introuvable**.). Strikingly, the only gene with a known function in terrestrial isopods encodes the androgenic gland hormone (AGH), which has a masculinizing effect and is involved in male sexual differentiation (Charniaux-Cotton, 1962; Martin *et al*., 1999; Suzuki, 1999). Non-synonymous causal SNPs affect five genes (Table 1), including one showing homology with *adam12*, a gene that may be linked to a receptor involved in sex determination in terrestrial isopods (see Discussion). None of the informative indels found in parents from both families were in the coding sequences of genes of the *M* locus. Four indels were localized in non-coding exons (UTRs), 78 in introns, and 11 at a distance < 1 kb from a protein-coding gene.

**Table 1.** Annotated genes in contigs that have not recombined with the *M* gene during crosses.

| Contig | Start | End | Strand | Putative product<sup>1</sup> | Non-synonymous SNPs<sup>2</sup> |
| --- | --- | --- | --- | --- | --- |
| Contig12786 | 28054 | 28897 | + | Hypothetical protein |  |
| Contig13319 | 8129 | 14688 | + | Lipoyltransferase/lipoate-protein ligase |  |
| Contig13319 | 18720 | 23796 | - | Dynein light chain, type 1/2 |  |
| Contig16348 | 21926 | 27485 | + | Glucose-methanol-choline oxidoreductase |  |
| Contig16348 | 28971 | 31378 | + | Glucose-methanol-choline oxidoreductase |  |
| Contig18030 | 4387 | 13386 | + | Protein Tramtrack |  |
| Contig18030 | 15988 | 16766 | - | Protein Bric a brac |  |
| Contig27690 | 17337 | 31848 | + | Adenylyl cyclase |  |
| Contig27690 | 43094 | 52696 | - | Peroxisomal ATPase | 1650: AAT->AAG N->K<br>1429: ACT->TCT T->S<br>1343: AGT->ATT S->I |
| Contig3268 | 81 | 6241 | - | Glutamine-tRNA ligase |  |
| Contig33327 | 22093 | 38297 | + | Hypothetical protein |  |
| Contig33327 | 40660 | 66415 | - | Lon protease |  |
| Contig35989 | 46505 | 52748 | - | Cilia- and flagella-associated protein 298 |  |
| Contig35989 | 58307 | 76688 | + | Nucleoside diphosphate kinase-like domain |  |
| Contig37362 | 8608 | 9165 | + | Hypothetical protein |  |
| Contig37362 | 41438 | 43179 | - | C-type lectin-like/link domain |  |
| Contig7365 | 12047 | 21034 | + | Glucose-methanol-choline oxidoreductase |  |
| Contig7365 | 20297 | 26887 | + | Glucose-methanol-choline oxidoreductase |  |
| Contig7365 | 43566 | 60107 | + | Broad-Complex, Tramtrack and Bric a brac (BTB) domain |  |
| Contig7365 | 73480 | 79840 | + | Chromatin modification-related protein Eaf7/MRGBP family |  |
| Contig7365 | 84129 | 86762 | - | Serine/threonine-protein kinase | 14: AGT->AAT S->N |
| scaffold_2521 | 37285 | 71933 | + | Arginine kinase |  |
| scaffold_2521 | 79282 | 88792 | - | Longitudinals lacking protein-like |  |
| scaffold_2521 | 97038 | 113787 | + | Protein tramtrack |  |
| scaffold_682 | 6 | 824 | - | Hypothetical protein |  |
| scaffold_682 | 23070 | 37974 | + | Ubiquitin thioesterase |  |
| scaffold_682 | 43871 | 61229 | + | T-complex protein 1 |  |
| scaffold_682 | 62507 | 63373 | - | Peroxisome biogenesis factor 2 |  |
| scaffold_682 | 70496 | 75117 | - | Androgenic gland hormone (AGH) |  |
| scaffold_682 | 87890 | 94749 | + | TIP41-like protein |  |
| scaffold_682 | 94809 | 99248 | - | RING-box E3 Ubiquitin Ligase |  |
| scaffold_682 | 107970 | 122401 | + | Protein-associating with the carboxyl-terminal domain of ezrin | 835: GTA->TTA V->L<br>2074: AAT->GGT N->G |
| scaffold_682 | 129143 | 142485 | - | 3'-5' ssDNA/RNA exonuclease TatD-like |  |
| scaffold_682 | 154273 | 224311 | - | Disintegrin and metalloproteinase domain-containing protein ADAM12 | 1231: CGT->AGT R->S |

<sup>1</sup>Only AGH is functionally characterized in *A. vulgare*. <sup>2</sup>The column indicates the position of nonsynonymous causal SNPs (see text) in gene coordinates (1-based) followed by the change in codon and the change in amino-acid.

**Figure 3.**
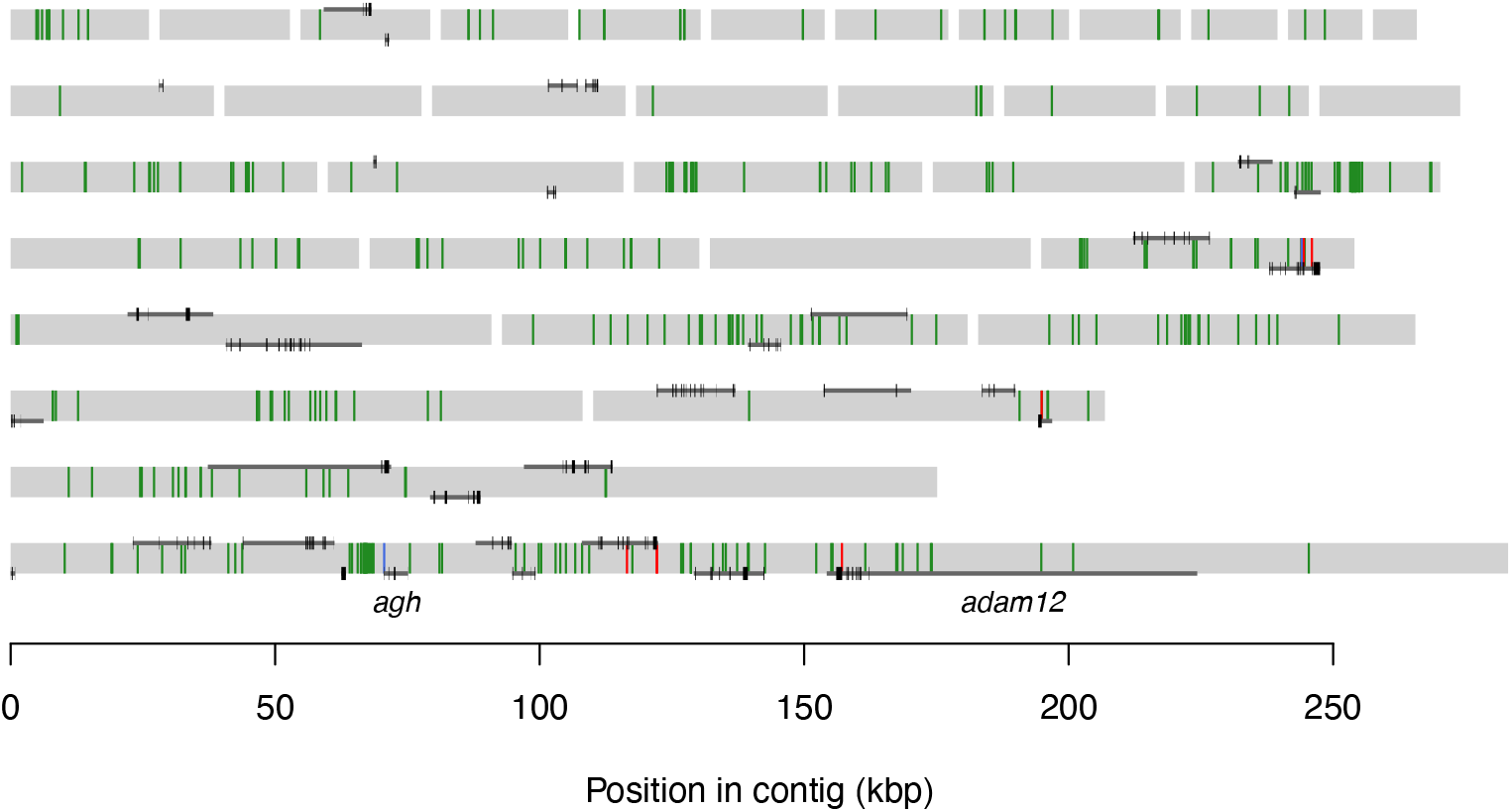
*Armadillidium vulgare* contigs that have not recombined with the *M* gene during crosses. Contigs are ordered by size from right to left and top to bottom and represented by grey rectangles. Dark grey segments represent predicted genes. Genes oriented form left to right are drawn on the top edge of contigs, otherwise they are drawn on the bottom edge. Thicker black segments within genes represent coding exons. Causal SNPs are drawn as vertical segments in the contigs (SNPs outside coding sequences are draw in green, synonymous SNPs in blue and non-synonymous SNPs in red). The *agh* and *adam12* genes are discussed in the text.

## Discussion

The genome-wide SNP analysis of daughters and sons of two *A. vulgare* families allowed narrowing down the *M* gene, a locus with a dominant masculinizing allele, to a ~1.9-Mb genetic region. Of the 34 annotated genes present in this region, the androgenic gland hormone gene (*agh*) is a prime candidate for being directly affected by the *M/m* polymorphism, given its crucial role in the male sexual differentiation cascade (Charniaux-Cotton, 1962; Martin *et al*., 1999; Suzuki, 1999; Herran *et al*., 2020). This hypothesis is supported by the fact that targeted injection of AGH at specific developmental stages induces sex reversal from female to male (Suzuki, 1999). Furthermore, when AGH is administered at a specific stages of female development (stages 6 and 7), it results in the development of males with female genital apertures (Suzuki and Yamasaki, 1995), as usually observed for males carrying the *M* gene. While no causal SNP appears to change the coding sequence of *agh*, other variants could *cis*-regulate *agh* expression. A change in the expression timeframe could explain the appearance of female genital apertures in males.

Another candidate gene carries a non-synonymous causal SNP and produces a protein identified as ADAM12 (Table 1). This protein is known to interact with the insulin-like growth factor-binding protein 3 (IGFBP-3) gene family in mammals (Loechel *et al*., 2000). In crustaceans, this gene family plays a central role in the AGH regulatory cascade (Rosen *et al*., 2013). Recent studies (Herran, 2018; Herran *et al*., 2019) have shown that in *A. vulgare*, IGFBPs exert an influence on sexual differentiation. However, it is not known whether the ADAM12 protein is involved in the male sexual differentiation cascade. If this protein is involved in regulating AGH, it is likely that its expression predominates in tissues associated with this hormone, such as the testes and androgen glands. This hypothesis, as well as the effect of the *M/m* polymorphism on *agh* and *adam12*, can be tested by comparing their transcription dynamics between *M/m* and *m/m* males during sexual differentiation. Comparing whole transcriptome data between these genotypes would allow characterizing the effect of the *M* allele on the expression of any gene, and to better understand its mode of action.

A complementary approach to narrow down the set of candidate genes affected by the *M/m* variation would be to refine the genomic region associated with sex, via a genome-wide association study (Kijas *et al*., 2018; Gabián *et al*., 2019; Wang *et al*., 2022). This approach would first require identifying natural populations where the *M* allele is present, for instance those including males carrying the *f* element or the W allele. Analysis of natural populations could also reveal traces of selective sweeps affecting the *M* locus, indicating that the *M* allele was recently selected possibly in response to a deficit of males induced by non-mendelian feminizing elements. As no increase in the density of heterozygous SNPs in males was observed close to the *M* gene, there is no indication that the *M* and *m* alleles have been diverging for a long period. Accordingly, the absence of homology between contigs linked to the *M* gene and those from the XY locus of *A. nasatum* indicates that these two XY-type loci emerged independently. These results do not contradict the hypothesis that the *M* allele has been selected recently in response to deficits of males induced by the *f* element (Rigaud *et al*., 1997), which has recently emerged (Leclercq *et al*., 2016). The apparent recent emergence of the *M* allele also reinforces the hypothesis of dynamic evolution of chromosomal sex determinants in terrestrial isopods (Becking *et al*., 2017). Population genomics analyses of sex-determinants in these species can provide empirical tests of theoretical models of sex chromosome renewal in response to sex-ratio distortion (Kozielska *et al*., 2010).

## Data availability

Scripts used to analyze data and are available at https://github.com/jeanlain/M-gene-Avulgare. Sequencing data are deposited in the SRA database at accession numbers [pending].

## Acknowledgements

We thank Dr Bouziane Moumen for providing support to bioinformatic analyses and Alexandra Lafitte for helping to identify males with female genital apertures. This work was funded by Agence Nationale de la Recherche grants ANR-15-CE32-0006 (CytoSexDet) to R.C., ANR-20-CE02-0004 (SymChroSex) to J.P., ANR-21-CE02-0004 (RESIST) to R.C., and by intramural funds from the CNRS and the University of Poitiers.

## Supplementary file description

### File S1.xlsx

This Excel file contains information about each contig analyzed. For each family, the posteriori probability of absence of recombination with the M gene is given, together with the posterior probability of the most likely haplotype frequency, and the read counts. Other columns include the contig name, its length, its inferred genetic distance to the SDR (*n*_rec_), whether it is assigned to sex chromosomes, and the heterozygous SNP density on fathers.

## Supplementary material

**Table S1.**
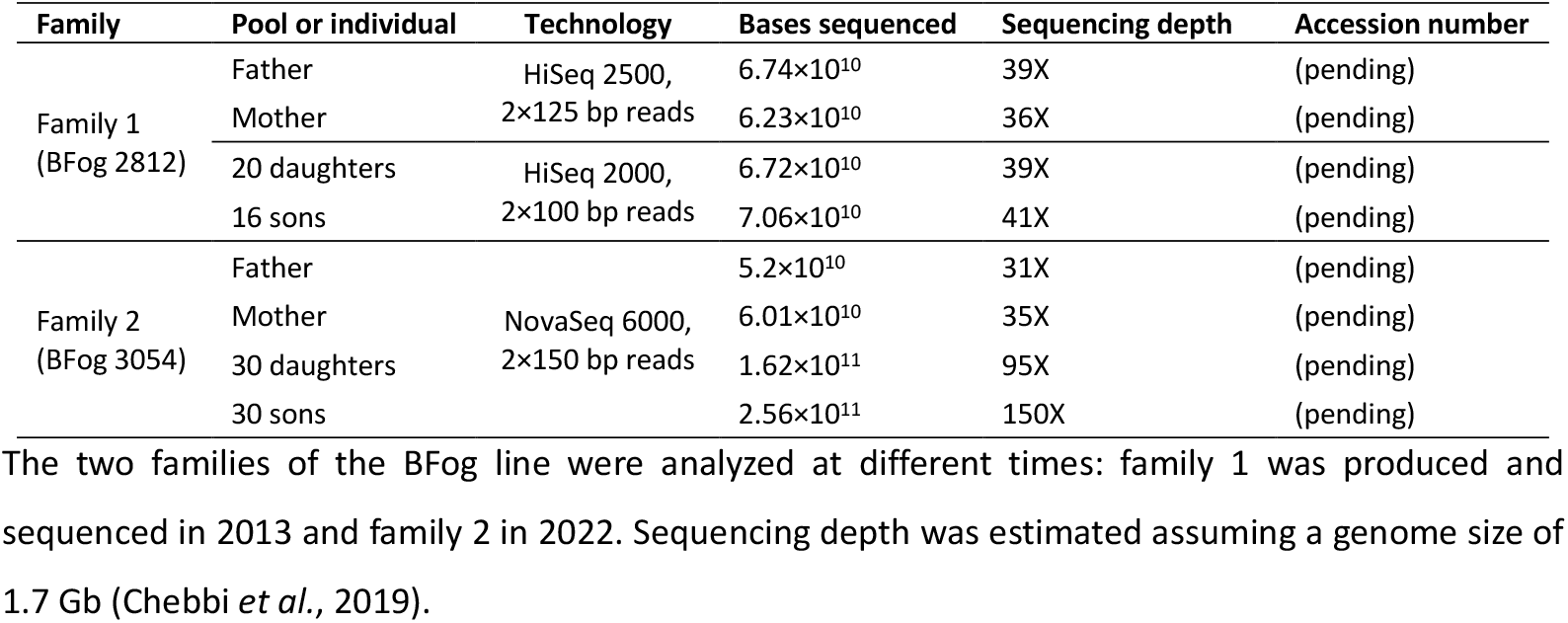
Sequencing statistics for the two families carrying the *M* allele.

**Table S2.**
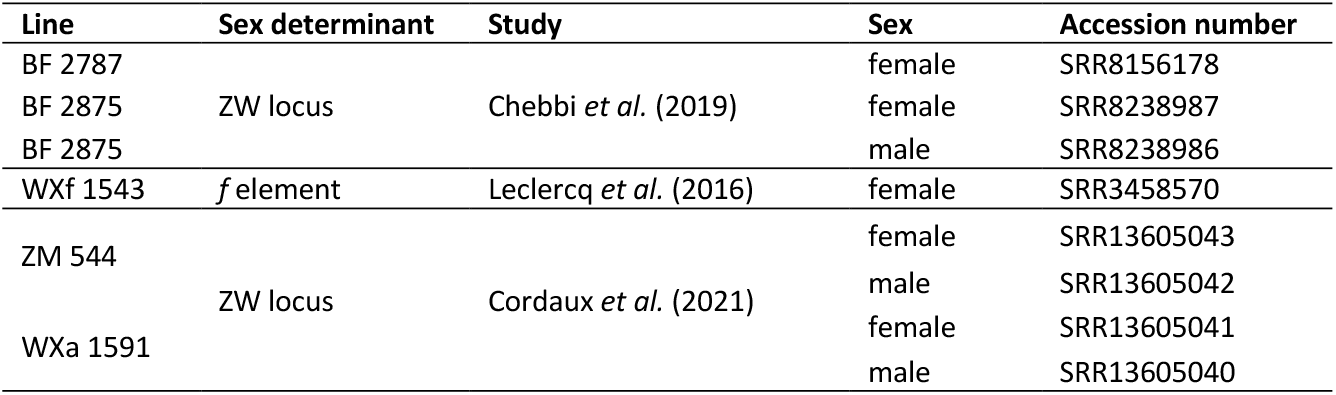
Sequenced individuals from lines not carrying the *M* allele.

### Presence of the W allele in analyzed families

Two primers were designed to amplify a target sequence linked to the W allele in contig 20397 (Cordaux *et al*., 2021) of the *A. vulgare* reference genome (Chebbi *et al*., 2019): 5’ATCTGTATGATTGGCCCATCCC3’ and 5’AACCTTTCGTAAGCGCCAAG3’. These primers were used in a quantitative PCR (qPCR) assay to estimate the dose of the W allele relative to a single-copy reference nuclear gene encoding the mitochondrial t-RNA leucine ligase (Durand *et al*., 2023). Expected relative doses are 0, 0.5 or 1, respectively indicating that the genotype is ZZ, ZW or WW. The qPCR mix used the Roche LightCycler 480 SYBR Green I Master in a 10 *μ*L total volume with 1 *μ*L of DNA template. The qPCR took place in a LightCycler 480 Instrument II with a denaturation phase at 94°C for ten minutes, followed by 45 cycles consisting of a denaturation step at 94°C for ten seconds, a 10-second hybridization phase at 60°C and then an elongation phase at 72°C for 20 seconds. Each DNA sample was amplified twice. The relative doses of the W allele were estimated by the Roche software using the second derivative method.

The qPCR assay confirmed the presence of the W allele in parents and offsprings of the second BFog family (**Figure**). Specifically, the mother was homozygous (relative dose ≈ 1) and the father heterozygous for this allele (relative dose ≈ 0.5). Expectedly, doses indicate the presence of homozygous and heterozygous descendants in similar proportions in both sexes. These results are consistent with the presence of the *M* allele in the father and in sons, as this allele can masculinize individuals that carry the W allele.

**Figure S1.**
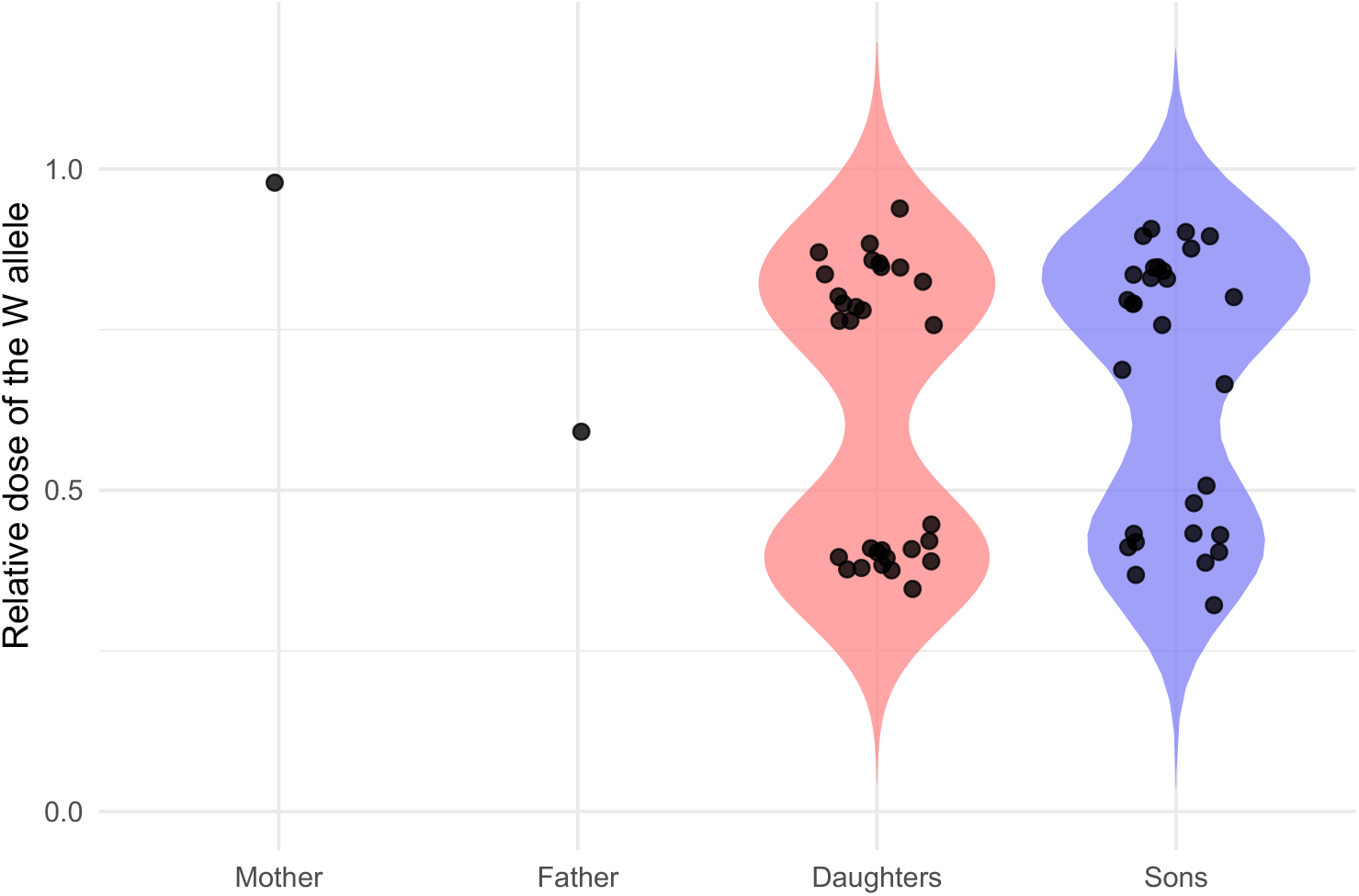
Relative dose of the W allele in parents and descendants of the second family of the BFog lineage, as estimated by quantitative PCR.

### Selection of informative SNPs

The analysis of SNPs to estimate the genetic distance between a contig and the *M* gene is essentially the same as that performed by Cordaux *et al*., (2021), which delineated the ZW locus of the same species, using the same reference genome. The pipeline was only slightly modified, mainly to account for male heterogamety rather than female heterogamety.

Informative SNPs were identified in each family as those meeting the following criteria. The posterior probability of the father being heterozygous and the mother homozygous, as output by ANGSD, must both exceed 0.95. Second, the sequencing depth must exceed 5 reads in each father, and 15 reads in each mother, and be less than the 95^th^ percentile of sequencing depth calculated over all SNPs in each parent. Third, the number of read pairs sequenced from the progeny and aligning on the SNP (hereafter “F1 reads”) must be ≥1 and be less than its 95^th^ percentile measured over all SNPs. Fourth, the paternal and alternative alleles must be carried by ≥75% of F1 reads. Fifth, the paternal allele of a SNP must have a frequency among F1 reads that does not significantly exceed 0.5, as determined via a one-sided binomial test with a p-value threshold of 0.05. Finally, we excluded any SNP for which the posterior probability of the most likely paternal allele frequency was much lower than most other SNPs of the same contig. This procedure is detailed in Cordaux *et al*., (2021) and was designed to mitigate the influence of read mapping errors (causing spurious SNPs) on the estimate of the paternal allele frequency at the contig level. Based on the selected SNPs, estimation of *n*_rec_ (see main text) for each contig in each F1 pool followed the principle described in Cordaux *et al*., (2021).

### Identification of causal SNPs

In contigs that were not found to have recombined with the *M* gene during crosses, “causal” SNPs were identified as informative SNPs for which both fathers and mothers respectively had the same genotype and for which the paternal allele frequency was higher in sons than in daughters (considering that for selected contigs, this frequency is either 0 or 0.5). This last criterion indicates that the paternal allele is linked to the *M* allele and that sons are heterozygous.

We also checked that supposed *m/m* individuals sequenced in previous studies (Table S2) lacked the paternal allele detected in individuals carrying the *M* allele. For each individual, aligned reads carrying each base of each SNP were counted by scanning the alignment (.bam) file with the mpileup function of samtools (Li *et al*., 2009), ignoring reads with mapping quality <20 (argument -q20). Defining *X* as the number of reads carrying a SNP allele, *x* its value for the SNP under scrutiny, and *n* the number of reads aligned on the SNP, the SNP was excluded if *P*(*X* ≤ *x*) > 5% [assuming that *X* followed *Binomial*(*n*, 0.5)], if *n* ≥ 5 and if the two alleles found in the BFog parents at that SNP were carried by at least *n*×0.9 reads.

### Estimation of SNP density in fathers

To compute the density of heterozygous SNPs in each father, we first selected SNPs with a probability of being heterozygous >99%, and a sequencing depth >5 and lower than its 95^th^ percentile over all SNPs. To estimate SNP density, we counted the number of positions where sequencing depth fulfilled these two conditions, for each contig, via the “depth” function of samtools on the alignment file used for SNP calling. To try to reproduce criteria used by ANGSD, reads with mapping quality lower than 1 were excluded (argument -Q 1) and bases with PHRED scores lower than 10 were ignored (argument - q 10) in sequencing depth computations. The density of heterozygous SNP per contig was computed as the number of selected SNPs in a contig divided by the number of positions meeting requirements on sequencing depth, combining both fathers.

